# Nanodiscs-based proteomics identify Caj1 as an Hsp40 with affinity for phosphatidic acid lipids

**DOI:** 10.1101/2021.06.18.448950

**Authors:** Xiao X. Zhang, John William Young, Leonard J. Foster, Franck Duong

**Affiliations:** Department of Biochemistry and Molecular Biology, Faculty of Medicine, Life Sciences Institute, University of British Columbia, Vancouver, British Columbia, Canada; Michael Smith Laboratory, University of British Columbia, Vancouver, British Columbia, Canada

## Abstract

Many soluble proteins interact with membranes to perform important biological functions, including signal transduction, regulation, transport, trafficking and biogenesis. Despite their importance, these protein-membrane interactions are difficult to characterize due to their often-transient nature as well as phospholipids’ poor solubility in aqueous solution. Here, we employ nanodiscs – small, water-soluble patches of lipid bilayer encircled with amphipathic scaffold proteins – along with quantitative proteomics to identify lipid-binding proteins in *S. cerevisiae*. Using nanodiscs reconstituted with yeast total lipid extracts or only phosphatidylethanolamine (PE-nanodiscs), we capture several known membrane-interacting proteins, including the Rab GTPases Sec4 and Ypt1, which play key roles in vesicle trafficking. Utilizing PE-nanodiscs enriched with phosphatidic acid (PEPA-nanodiscs), we specifically capture a member of the Hsp40/J-protein family, Caj1, whose function has recently been linked to membrane protein quality control. We show that Caj1 interaction with liposomes containing PA is modulated by pH and PE lipids, and depends on two patches of positively charged residues near the C-terminus of the protein. The protein Caj1 is the first example of an Hsp40/J-domain protein with affinity for membranes and phosphatidic acid lipid specificity. These findings highlight the utility of the nanodisc system to identify and characterize protein-lipid interactions that may not be evident using other methods.

## INTRODUCTION

Acidic phospholipids such as phosphatidylinositols (PIs), phosphatidic acid (PA) and phosphatidylserine (PS) are relatively minor constituents of eukaryotic membranes, yet these lipids play critical roles in recruiting protein effectors to the plasma and organellar membranes ^1,2^. These membrane-bound effectors in turn regulate important signal transduction pathways, in addition to membrane biogenesis and protein trafficking processes ^1, 3–5^. Identification of the membrane-bound proteome is therefore central to the understanding of membrane biology. It is however experimentally challenging given the insoluble nature of the membrane and the overall transient nature of protein-lipid interactions ^3, 4^.

Isolation of lipid bound effectors to date has enabled identification of the main protein binding domains that interact with acidic phospholipids. Notable examples include the Plextrin Homology (PH) and PhoX Homology (PX) domains, which recognize different phosphorylation states of PI lipids, as well as the C2 domain which interacts with PS ^1^. No such domain, however, has been discovered for proteins that associate to PA ^3, 5, 6^. The only shared feature between PA-binding proteins appears to be clusters of positively charged amino acids, which may be important for forming electrostatic interactions with negatively charged lipids ^3, 5–7^. There is however no sequence conservation for these clusters that explains lipid specificity, rendering bioinformatic identification of novel PA-binding proteins difficult ^3, 4, 8^. Consequently, the number of known PA-binding proteins is relatively low compared to those known to associate with the acidic lipids PIs and PS ^1, 8, 9^.

Most PA-binding proteins have been identified so far using PA lipid molecules covalently attached to polymer beads, followed by affinity pull-down and mass spectrometry analysis ^9^. The major drawback of this method, however, is that it does not faithfully mimic a native membrane bilayer. Thus, potential interactors which require a proper lipid bilayer surface for high-affinity binding may escape detection.

Numerous membrane mimetics have been developed in the recent years to try to mimic the native membrane environment in aqueous solution ^10–13^. Our previous work using nanodiscs and peptidiscs combined with SILAC labeling and quantitative proteomics has demonstrated the utility of these systems to identify and characterize protein-protein interactions taking place at the membrane interface ^14, 15^. This present study develops a similar proteomics strategy to identify novel PA-interacting proteins in the yeast *S. cerevisiae* (Figure 1).

**Figure 1:**
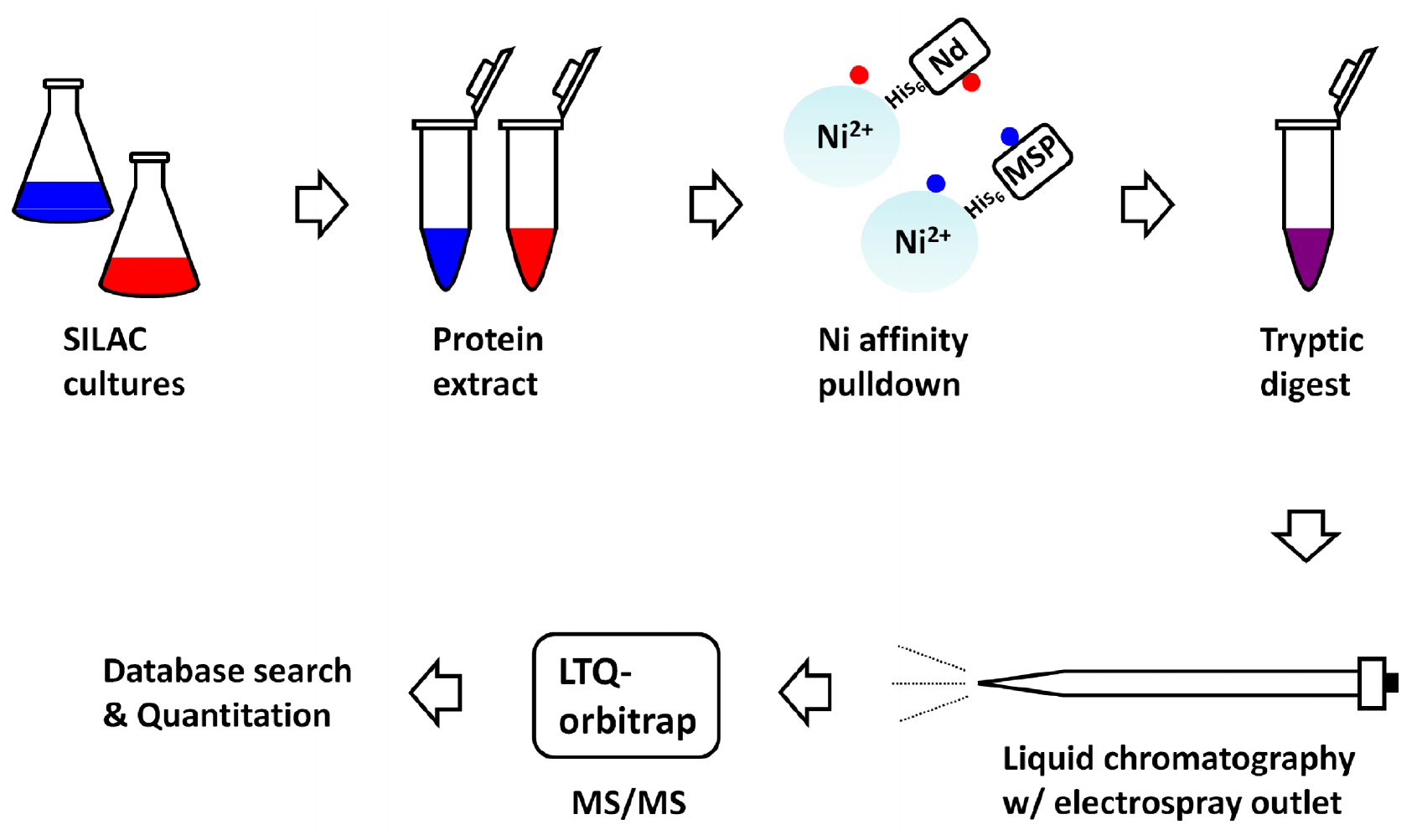
Overview of the nanodisc-SILAC proteomics workflow. SILAC-labeled “Heavy” (red) and non-labeled “Light” (blue) yeast soluble protein extracts were subjected to Nickel affinity pulldown using either lipid nanodiscs (“Heavy” extract) or MSPs (“Light” extract) as bait. After pooling the samples, the eluate was digested with Trypsin and desalted by STAGE tipping before analysis by LC-MS/MS.

To benchmark our method, we first used nanodiscs made with total yeast lipid extracts. As expected, these nanodiscs capture several known lipid-binding proteins, including the Rab GTPases Sec4 and Ypt1. Since phosphatidic acid is only a minor constituent of the yeast membrane, we next constructed PA-enriched nanodiscs. In that case, we identify the J-domain containing protein Caj1 as a high-affinity interactor of phosphatidic acid. We confirm this interaction using liposomes and we show that Caj1 interacts specifically with PA lipids due to two patches of positively charged residues located in the C-terminal domain of Caj1. We also find that the interaction of Caj1 with PA lipids is pH dependent. These results are striking in the context of recent *in vivo* data, which suggests that Caj1 may be involved in membrane quality control ^16, 17^. While it is not currently known how Caj1 performs its function, it has been reported that Caj1 is partially localized to the yeast plasma membrane ^17^. Our observation that Caj1 displays a high affinity for lipid bilayers containing phosphatidic acid provides insights into how this Hsp40 may be specifically targeted to the cell membrane.

## MATERIALS AND METHODS

### Reagents

Phospholipids were obtained from Avanti Polar Lipids Inc. (Alabaster, AL). Yeast nitrogen base was purchased from Becton, Dickinson and Company (Sparks, MD). Complete supplement mixture minus arginine and lysine was obtained from Sunrise Science Products, Inc. (San Diego, CA). Isotopically labelled lysine and arginine were from Cambridge Isotope Laboratories (Andover, MA). Detergent n-dodecyl-β-d-maltoside (DDM) was purchased from Anatrace. The resin Ni^2+^-NTA chelating Sepharose was obtained from Qiagen. Tryptone, yeast extract, NaCl, imidazole, Tris-base, acrylamide 40%, bis-acrylamide 2% and TEMED were obtained from Bioshop Canada. All other chemicals were obtained from Fisher Scientific Canada.

### Preparation of SILAC-labeled cultures

The lysine and arginine auxotrophic strain, BY4742 *Δarg4*, was from the Loewen laboratory collection ^18^. The cells were first grown for 7 h in YPD media at 25°C, washed three times with sterile water and diluted to a final O.D._600_ of 0.007 in yeast nitrogen base minimal medium. For the “light” labeled culture, media contained 0.69 mg/ml complete supplement mixture minus arginine and lysine, 36 μg/ml of arginine, and 30 μg/ml of lysine. For the “heavy” SILAC labeled culture, ^13^C_6_ ^15^N_4_-arginine and ^13^C_6_ ^15^N_2_-lysine was added in place of arginine and lysine. The cultures were grown side-by-side for 14 h at 25 °C prior to harvesting. Cell pellets were re-suspended in TSG buffer (50 mM Tris-HCl pH 7.9, 50 mM sodium chloride, 10% w/v glycerol) and lysed in the presence of 10 mM protease inhibitor by glass beads (1.5 ml of beads per 1 ml of culture). Lysates were centrifuged at 7000 × *g* to remove cellular debris, then at 200,000 × *g* where the soluble protein fraction was obtained.

### Nanodisc preparation

The His-tagged membrane scaffold protein MSP1D1 (MSP) was recombinantly expressed and purified exactly as previously described ^19^. Phospholipids dissolved in chloroform were dried under a stream of nitrogen and left in a vacuum desiccator overnight. For nanodiscs containing mixtures of lipids, phospholipids were mixed in the desired molar ratio in chloroform before being dried down. Lipids were then resuspended in TSG buffer supplemented with 0.1% DDM by gentle pipetting. Nanodiscs were prepared as previously described with modifications ^20^. Briefly, purified MSP and phospholipids were mixed in a 1:20 molar ratio. To remove detergent and facilitate nanodisc formation, the mixture was incubated with BioBeads overnight at 4 °C. All discs were purified by size exclusion chromatography on a Superdex 200 10/300 column equilibrated in TSG buffer. Peak fractions were pooled, concentrated, and stored at −80 °C until use.

### Pulldown experiments and mass spectrometry

Affinity pull-down of the nanodiscs with SILAC extract was performed following Zhang *et al.* ^14^. Briefly, 10 μg nanodisc or MSP were bound onto 20 μL Nickel NTA agarose resin for 5 min at 25 °C in TSG buffer with gentle shaking. After a brief wash to remove unbound material, the resin bound with nanodisc was incubated for 30 min with 1 mg “heavy” SILAC labeled yeast cytosolic extract. As a control, 1 mg “light” cytosolic extract was incubated with the resin bound with MSP. Beads from both samples were then combined and washed 3 times with 1 mL TSG buffer supplemented with 50 mM Imidazole. Proteins were eluted in 100 μL TSG buffer supplemented with 600 mM Imidazole. Proteins were digested with Trypsin, STAGE-tipped and analyzed by mass spectrometry exactly as previously described ^14^. The yeast ORF database was used to identify peptides/proteins in Mascot. Protein ratios were quantified using Proteome Discoverer v1.2 (Thermo Fisher Scientific). Proteins identified with only a single unique peptide were not included in our analysis.

### Caj1 purification

The gene encoding Caj1 was amplified from yeast genomic DNA prepared from the strain BY4742 and cloned into a pET23a vector between the Nde1 and Xho1 restriction sites. Successful cloning was verified by DNA sequencing. Point mutations were introduced by site-directed mutagenesis and confirmed by DNA sequencing. pET23a Caj1 (wild-type and mutant variants) was transformed into *E. coli* C41 cells and grown with shaking at 37 °C in 1 L LB media supplemented with 100 μg/mL Ampicillin. Protein expression was induced with 0.5 mM IPTG at OD_600_ 0.4. Cells were grown a further 3 h before harvesting by centrifugation (5000 × g, 10 min). Cells were homogenized in cold TSG buffer supplemented with 1 mM PMSF and lysed by 3 passages through a Microfluidizer (15,000 psi). Unbroken cells were removed by centrifugation (5,000 × g, 10 min). The lysate was further subjected to ultracentrifugation (100,000 × g, 40 min) to remove membranes and large aggregates before being loaded onto a 5 mL SP-Sepharose column pre-equilibrated in TSG buffer on an AKTA FPLC. After loading the sample, the column was washed extensively with TSG buffer. Bound proteins were eluted with a 50 mL gradient from 50 mM – 1 M NaCl. All fractions were analyzed by 15% SDS-PAGE and Coomassie Blue staining to assess purity. Fractions containing the Caj1 protein were concentrated in a Centricon filter (10 kDa cutoff) and further purified by size-exclusion chromatography on a Superdex 200 10/300 column.

### Native PAGE and affinity purification experiments

Recipes and protocols for Native PAGE analysis of protein samples have been described in detail in Carlson *et. al.* (2018) ^21^. In the Caj1 titration binding experiments, the amount of nanodisc (nanodisc-PEPA or nanodisc-PE) was held constant at 1.5 μg across all conditions. Increasing amounts of Caj1 was added to the discs. All samples were mixed on ice for 30 min in a 20 μL volume before being loaded onto the gel. After electrophoresis, gel bands were visualized by Coomassie blue staining. Affinity purification experiments using nanodiscs and purified Caj1 were performed essentially as previously described ^22^. Briefly, 10 μg nanodisc (or MSP alone) were bound onto 20 μL Nickel NTA agarose resin for 5 min at 25 °C in TSG buffer with gentle shaking. After a brief wash to remove unbound material, the resin bound with nanodisc was incubated for 30 min with 20 μg purified Caj1 in a 1 mL volume. The resin was then washed 3 times with 1 mL TSG buffer. Proteins were eluted in 50 μL TSG buffer supplemented with 600 mM Imidazole and visualized by SDS-PAGE and Coomassie blue staining.

### Membrane sedimentation experiments

Phospholipids were dissolved in chloroform and mixed in the desired ratios. The lipid mixtures were then dried under a stream of nitrogen and overnight in a vacuum. The lipids were then resuspended in TSG buffer using a freeze-thaw method (flash frozen in liquid nitrogen, then rapidly thawed at 37 °C and vortexed). The freeze-thaw step was repeated five times to ensure uniformly size liposomes. Liposomes used in this study include: PC (100), PCPE (60:40), PCPEPA (90-50:0-40:10), PCPEmPA (50:40:10). All lipid ratios were calculated in moles. In 150 μl TSG buffer, 150 nmol of liposomes and 11.5 μg of wild-type or mutant variants of Caj1 were added. Mixtures were incubated for 20 min on ice followed by ultracentrifugation at 174,000 × *g* for 30 min at 4 °C. Pellets were washed once in TSG buffer then sedimented again by ultracentrifugation. The pellets were analyzed by SDS-PAGE.

## RESULTS

### Identification of phospholipid interacting proteins

To benchmark our nanodisc-SILAC experimental approach and demonstrate its effectiveness for isolating lipid-interacting proteins in yeast, we reconstituted yeast total phospholipid extract into nanodiscs as described in the Materials and Methods. Nanodiscs were immobilized on Ni-NTA resin and incubated with SILAC-labeled “heavy” yeast soluble protein extracts (Figure 1). To measure protein enrichment and to control for non-specific background contaminants, MSP was immobilized on Ni-NTA resin and incubated side-by-side with non-labeled “light” yeast soluble protein extracts. The two batches of resin were then pooled, and bound proteins were eluted in TSG buffer supplemented with 600 mM Imidazole (Figure 1). The eluate was digested with Trypsin and desalted using STAGE tips prior to LC-MS/MS analysis. To decrease the data variability, the mass spectrometry experiments were performed twice using two independent biological replicates.

A total of 51 proteins were present across both replicates and were identified with at least two unique peptides. These proteins were listed in descending order based on their average SILAC ratio (Supplemental File 1). As a graphic representation of the data, the average SILAC ratio for each protein was plotted as a function of its rank in the protein list (Figure 2A). Using this ranking method which report proteins highly enriched in the nanodisc pulldown relative to the MSP control, we identified 3 predominant proteins: the Rab GTPases Sec4 and Ypt1, and the fatty acid synthetase Faa1. These three soluble proteins are known to perform their function at the membrane, suggesting that our method is effective for capturing transient and peripherally bound interactors of phospholipids ^23, 24^.

**Figure 2:**
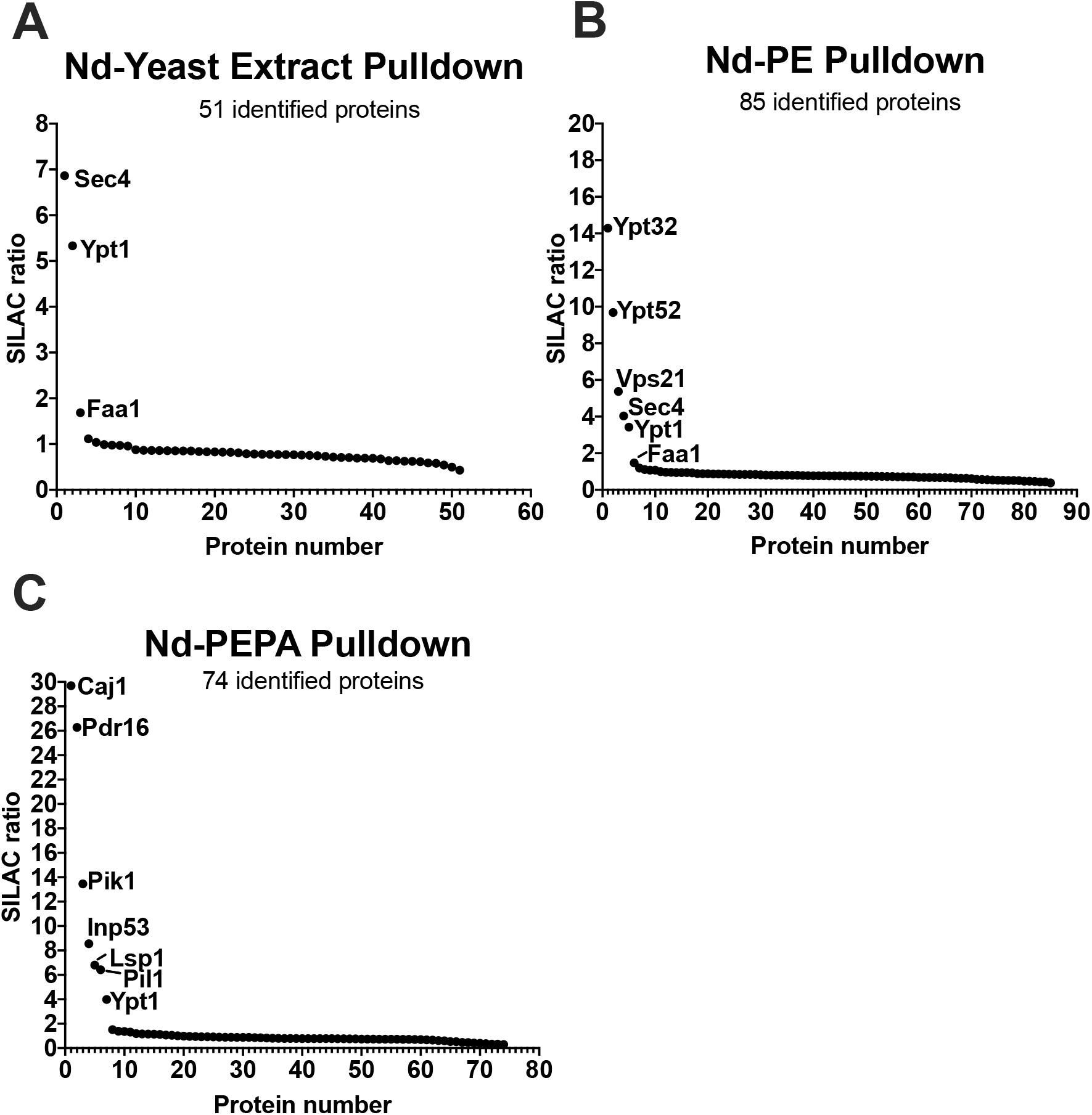
SILAC ratios for proteins identified in nanodisc pulldowns. **A)** SILAC ratios for proteins identified in our yeast extract nanodisc pulldown were plotted against their rank in the protein list. Only proteins with a SILAC ratio greater than 1 are labeled. Each point represents the average of 2 independent experiments. **B)** As in (A), but for our PE nanodisc pulldown. **C)** As in (A), but for our PEPA nanodisc pulldown.

To further determine if our method can be applied to identify lipid-specific interactors, we started using nanodiscs containing phosphatidylethanolamine (PE) only. Nanodiscs containing pure PE were employed because PE lipid is major structural membrane component across multiple different organelles in yeast ^25^. After incubation with yeast soluble extract and affinity pulldown, the mass spectrometry data was analyzed and plotted as described above (Supplemental File 1, Figure 2B). Out of a total of 85 identified proteins, we observe five proteins enriched with SILAC ratios greater than 1. As seen above, the predominantly identified proteins using this workflow are member of the Rab GTPases family, namely Ypt32, Ypt52, Sec4, Vps21 and Ypt1. These Rab GTPases play key roles in membrane trafficking, such as endocytosis, exocytosis, and vesicle secretion. These Rab GTPases have a common mechanism for membrane localization via their carboxyl-terminal prenylation ^23, 24^. Collectively, these results show that our method is suitable for the identification of membrane interacting proteins.

### Identification of phosphatidic acid interacting proteins

Next, we applied the same nanodisc-SILAC workflow to search for interactors of the anionic phosphatidic acid using lipid nanodiscs containing mixture of PE and PA at a 50:50 molar ratio. Strikingly, after incubation with yeast soluble extract and affinity pulldown, the most highly enriched protein identified in our mass spectrometry data list is the protein Caj1 (Supplemental File 1, Figure 2C). Caj1 is Hsp40 heat shock protein belonging to the *E. coli* DnaJ family. While its exact biological role in the cell context remains unclear, recent *in vivo* work has suggested that Caj1 may be involved in membrane protein quality control by an as-yet uncharacterized mechanism ^16, 17, 26^. Besides Caj1, we also identify the phosphatidylinositol (PI)-interacting proteins Pdr16, Pik1 and Inp53. Pdr16 is annotated as a phosphatidylinositol transfer protein; Pik1 is a phosphatidylinositol kinase, and Inp53 is a phosphatidylinositol phosphatase. Our observation that proteins involved with PI modification interact with PA is not entirely surprising because previous work has shown that active site in phosphatidylinositol kinases also display affinity for PA ^8^.

### Recombinant expression and purification of Caj1

To validate our proteomic observation that Caj1 interacts with PA, we first cloned the gene encoding Caj1 from *S. cerevisiae* genomic DNA into a bacterial protein expression vector to enable recombinant expression in *E. coli*. Caj1 was purified from the clarified cell lysate over SP-Sepharose cation exchange resin as described in the Materials and Methods. SDS-PAGE analysis of the purified protein after cation exchange chromatography reveals a prominent band at ~ 45 kDa, the expected molecular mass of Caj1 (Figure 3A). Surprisingly, we observe two smaller co-purifying bands at ~ 30 kDa (indicated by *) and 10 kDa (indicated by †) (Figure 3A). To identify these co-purifying proteins, we excised the bands from the gel and analyzed them by mass spectrometry. Our analysis reveals both bands contain peptides derived from Caj1, indicating the bands represent proteolytic fragments of Caj1. To increase the purity of our material, we further purified Caj1 by size-exclusion chromatography (Figure 3B). SDS-PAGE analysis of the peak protein fractions reveals that the ~ 30 kDa and ~ 10 kDa proteolytic fragments co-migrate with intact Caj1 during the chromatography.

**Figure 3.**
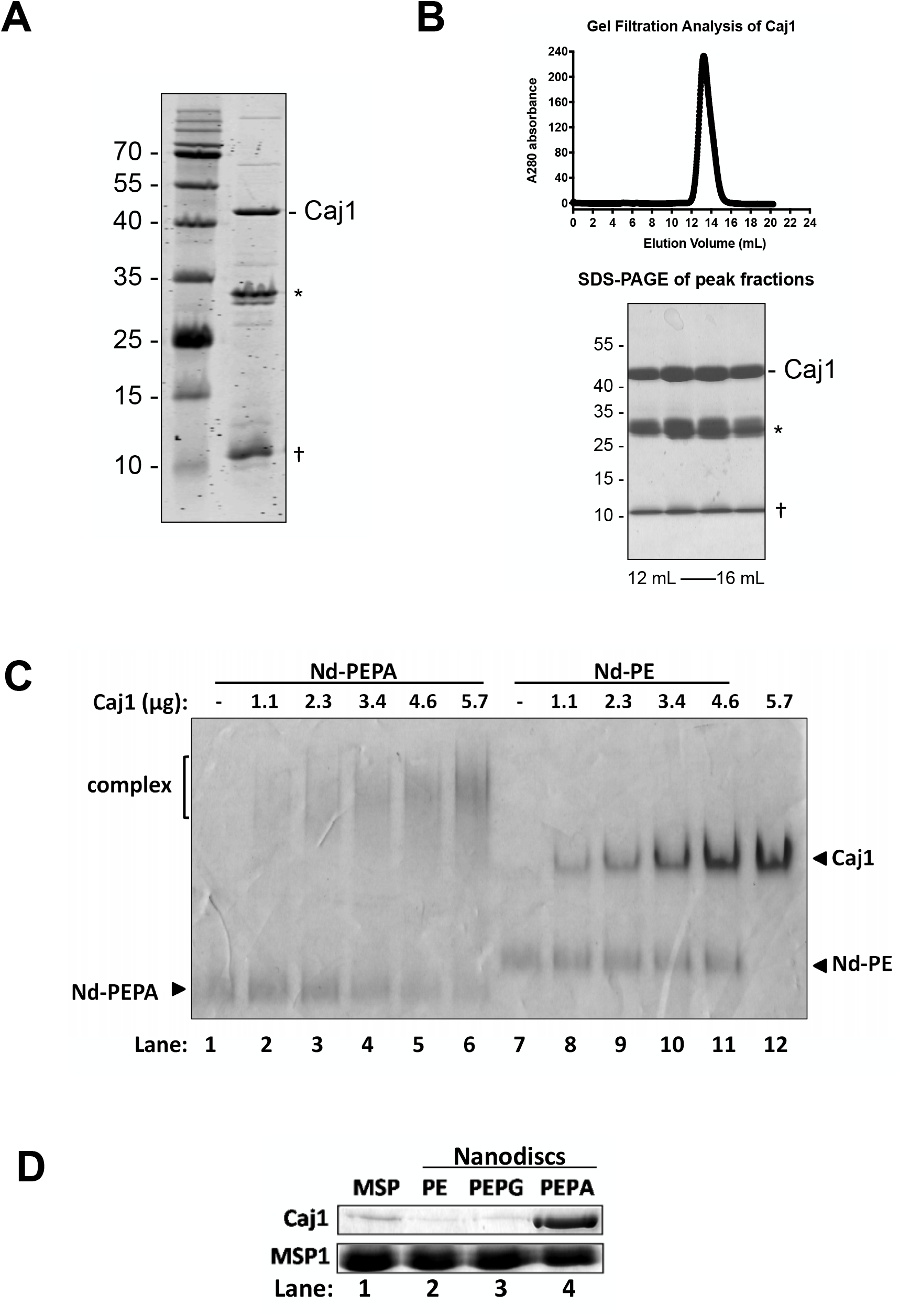
Purification of Caj1 and interaction with PA lipids. **A.** Representative SDS-PAGE of Caj1 following cation exchange chromatography. The position of intact Caj1 is indicated. The position of the ~30 kDa and ~10 kDa proteolytic fragments are indicated by * and †, respectively. **B**. Caj1 from (A) was further purified by size-exclusion chromatography on a Superdex 200 10/300 column. UV absorbance as a function of elution volume was plotted (upper panel). Fractions under the main peak (12 mL-16 mL) were further analyzed by SDS-PAGE (lower panel). **C.** Clear-native PAGE (4-12 %, pH 8.8) analysis of Caj1-PA interaction using Nanodisc. Indicated amount of Caj1 was incubated with 1.5 μg of Nanodisc containing either PEPA (50:50 molar ratio) or PE. Bands on all gels were detected by Coomassie blue staining. **D.** Affinity pull-down of Caj1 with Nanodiscs. Nanodiscs-PE, PEPG (50:50 molar ratio), and PEPA (50:50 molar ratio) were immobilized onto Ni-NTA beads. MSP alone was immobilized on resin side-by-side as a control. A solution of Caj1 in TSG buffer was passed over the resin. After a brief wash to remove unbound material, proteins were eluted from the resin with TSG buffer plus 600 mM imidazole. Proteins were analyzed on SDS-PAGE followed by Coomassie blue staining.

### Verification of Caj1-PA interaction using nanodiscs

To verify the interaction between Caj1 and PA, we performed electrophoretic mobility shift assays (EMSA) on non-denaturing native PAGE (Figure 3C). Caj1 was titrated against a fixed amount of nanodisc-PEPA. The same titration was performed side-by-side using nanodisc-PE to ensure any observed interactions are specific to PA. Caj1 alone migrates as a single, well-resolved band on native PAGE (Figure 3C, lane 12). Nanodisc-PEPA and nanodisc-PE also form discrete bands on the gel (Figure 3C, lanes 1 and 7). As progressively increasing amounts of purified Caj1 are added to nanodisc-PEPA, we observe the formation of a higher molecular weight species in the gel, indicating that Caj1 is binding onto the nanodisc (Figure 3C, lanes 2-6). No such interaction is observed when the same titration is performed using nanodisc-PE (Figure 3C, lanes 8-11), indicating that the observed interaction is specific to PA.

We also performed affinity pulldown experiments using lipid nanodiscs and Caj1 (Figure 3D). Nanodiscs or MSP alone were immobilized on Ni-NTA resin. A solution containing purified Caj1 was then passed over the resin. After a brief wash with TSG buffer, bound proteins were eluted with TSG supplemented with 600 mM imidazole and visualized on SDS-PAGE. In agreement with our EMSA results, Caj1 binds with nanodisc-PEPA (Figure 3D, lane 4) but not with MSP alone or with nanodisc-PE (Figure 3D, lanes 1 and 2). To confirm that Caj1 interacts with PA specifically, rather than with acidic lipids in general, we also tested whether Caj1 interacts with nanodiscs containing phosphatidylglycerol (PG). Caj1 does not co-purify with nanodisc-PEPG (Figure 3D, lane 3), showing that the Caj1-phospholipid interaction is dependent on PA.

### Verification of Caj1-PA interaction using liposomes

To complement our nanodisc binding assays, we assessed the ability of Caj1 to interact with PA in liposomes. Phospholipids mixed in defined ratios were resuspended in buffer and subjected to multiple freeze-thaw cycles to form liposomes as described in the Materials and Methods. Liposomes were incubated with purified Caj1 before being pelleted by ultracentrifugation. Pellets were resuspended in buffer and analyzed by SDS-PAGE to visualize bound proteins (Figure 4A). As expected, Caj1 does not sediment with liposomes containing only the major yeast structural lipid phosphatidylcholine (PC) (Figure 4A, lane 1), nor does it sediment with liposomes containing a 60:40 mixture of PC and PE (Figure 4A, lane 2).

**Figure 4.**
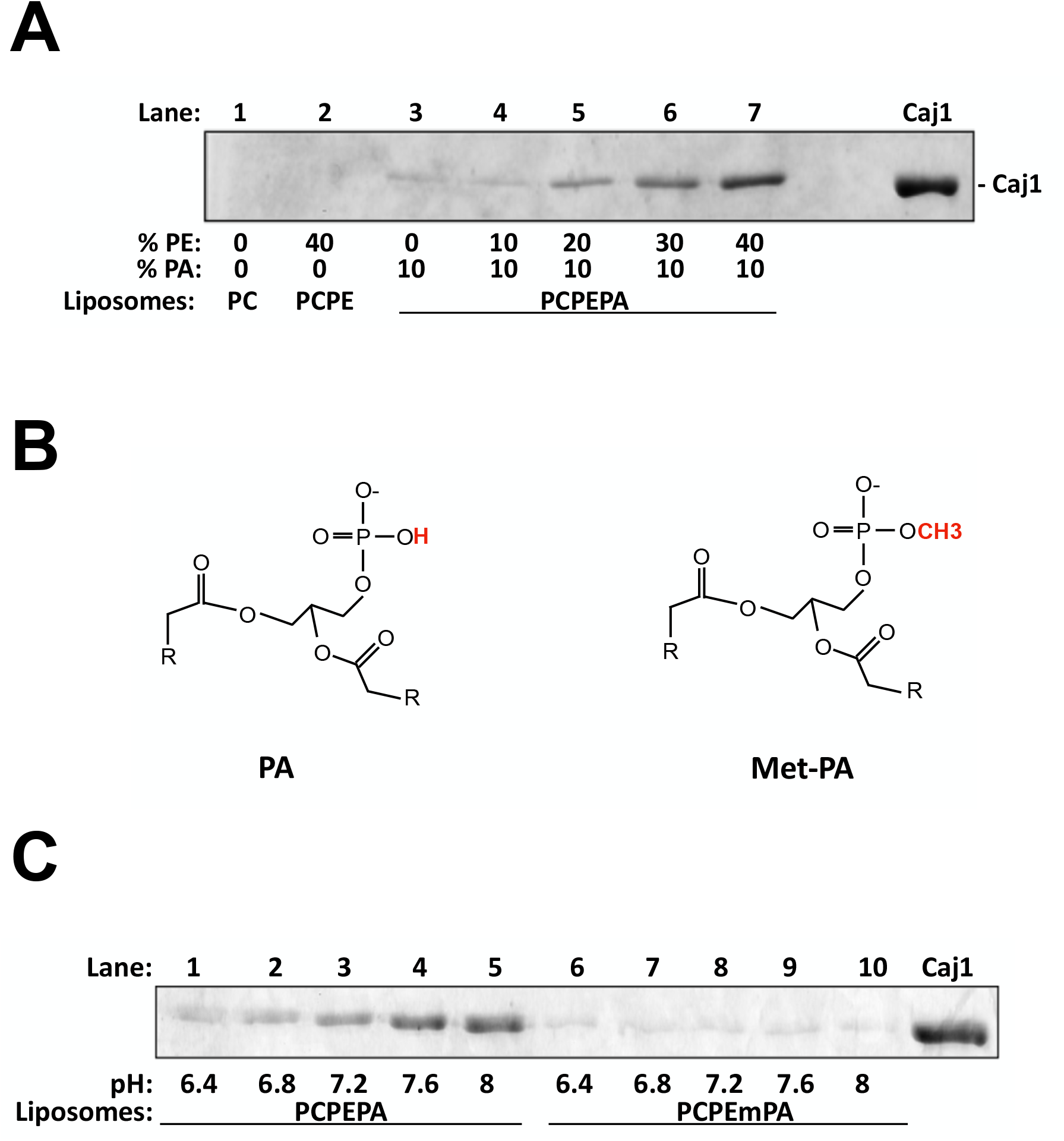
Caj1 interaction with PA in liposomes. **A.** Caj1 was incubated with liposomes containing phospholipids at the indicated molar ratios. Liposomes were sedimented by ultracentrifugation and resuspended in buffer. Bound proteins were visualized on SDS-PAGE followed by Coomassie blue staining. As a control, 2 μg Caj1 was loaded on the right-most gel lane (lane “Caj1”). **B.** Structures of phosphatidic acid (left panel) and the methylated derivative (Met-PA, right panel). The ionizable proton of PA (pK_a_ ~ 7) is highlighted in red. The non-ionizable methyl substitution on Met-PA is also highlighted. Note that “R” groups denote the lipid alkyl chains. **C.** Effect of pH and headgroup charge on the Caj1-PA interaction. Caj1 was incubated with PCPEPA (50:40:10 molar ratio) or PCPEmPA (50:40:10 molar ratio) liposomes in TSG buffer at the indicated pH. Bound proteins were visualized as described in (A).

Previous studies have shown using sedimentation assays that increasing the concentration of PE in liposomes containing PA leads to increased binding of PA-interacting proteins ^6, 27^. This could be because PE increases the negative charge on the PA headgroup from −1 to −2 via hydrogen bonding interactions ^6, 7^. Since interactions between PA and its protein binding partners are largely driven by electrostatics, this increase in charge should lead to greater binding. To test if this prior observation is applicable to Caj1, we fixed the concentration of PA in our liposomes at 10% (mol/mol) while the concentration of PE was titrated from 0-40% (Figure 4A, lanes 3-7). In line with previous reports, we observe greater sedimentation of Caj1 as the concentration of PE in the liposomes increases ^6, 27^.

Next, we examined the effect of pH on binding of Caj1 onto PA. The pK_a_ of the phosphomonoester headgroup of PA is within physiological range (pH ~7) ^5, 27^. Thus, slight changes in pH between pH ~6-8 should influence the overall charge of PA which may in turn affect Caj1 binding. To assess this possibility, we formed liposomes containing 50% PC, 40% PE and 10% PA. Liposomes were resuspended in buffer with pH varying between 6.4 – 8 before being incubated with purified Caj1 protein. Liposomes were sedimented and bound proteins analyzed as described above. In agreement with previous reports, there is greater sedimentation of Caj1 with the liposomes at higher pH (Figure 4C, lanes 1-5) ^27^.

To highlight the importance of the charge on the headgroup of PA for the Caj1-PA interaction, we also employed a non-titratable methylated PA derivative, termed Met-PA (Figure 4B). Due to the methyl group, changes in pH are not expected to affect the overall charge of Met-PA. We formed liposomes containing 50% PC, 40% PE and 10% Met-PA and performed sedimentation assays as described above (Figure 4C, lanes 6-10). Strikingly, we observe very little binding of Caj1 onto liposomes containing Met-PA, and there is no apparent effect of pH. Altogether, these results underscore the importance of the charge on the PA headgroup for the Caj1-PA interaction and highlight how this charge is influenced by slight changes in environmental pH.

### Characterization of potential PA binding site on Caj1

Next, we attempted to identify residues within Caj1 that may be responsible for mediating its interaction with PA. Since the N-terminal 80 residues of Caj1 constitute the conserved J-domain, we confined our search to the more variable C-terminal portion of the protein ^26, 28^. For the few other characterized PA-interacting proteins, PA recognition is mediated through short clusters of positively charged residues – often lysine or arginine ^3, 6, 9^. Examination of the sequence of Caj1 (https://www.uniprot.org/uniprot/P39101 – accessed April 23, 2021) reveals two double lysine motifs – KK 245-246 and KK 335-336. For clarity, we term KK 245-246 “Patch 1” and KK 335-336 “Patch 2” (Figure 5A). Analysis of the Caj1 sequence using the software DNA strider reveals that the entire protein has a slightly acidic pI of 5.79 ^29^. However, the regions surrounding “Patch 1” and “Patch 2” have a pI of 10.52 and 9.84 respectively (Fig. 5A). To determine whether the Patch 1 and/or Patch 2 residues are involved in mediating the Caj1-PA interaction, we generated double-alanine mutations, generating “Patch 1 Ala” and “Patch 2 Ala” Caj1. We also constructed a “Patch 1+2 Ala” double mutant. All mutant constructs behaved comparable to the wild-type protein during expression and purification.

**Figure 5.**
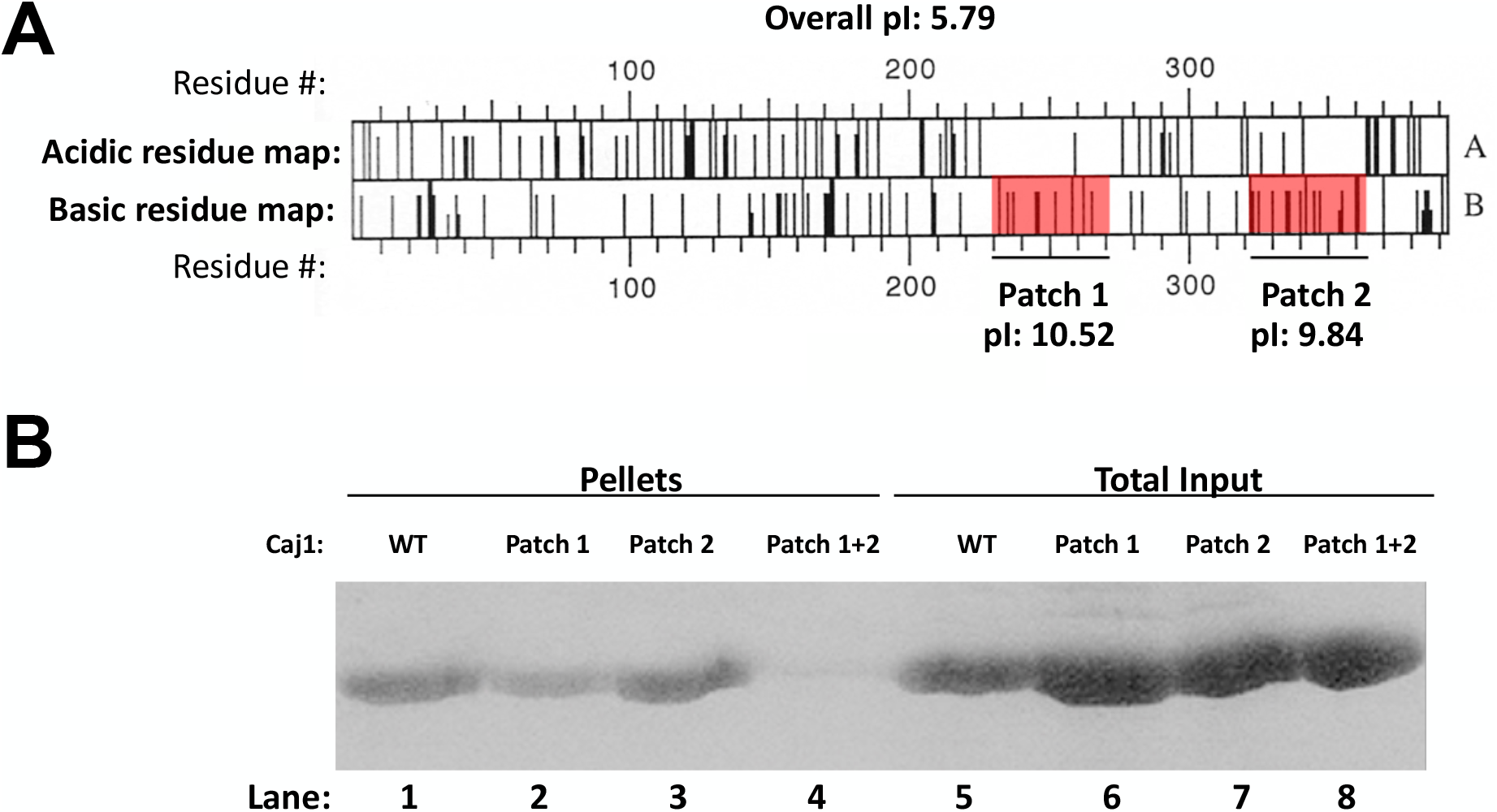
PA binding assay of Caj1 mutants. **(A)** Schematics of acidic (top panel) and basic regions (bottom panel) on Caj1. Black bars represent the acidic or basic residues. Highly basic regions towards the C-terminus are highlighted in red and indicated as “Patch 1” and “Patch 2” as appropriate. Map is generated from N-terminus to C-terminus, left to right, using the DNAStrider program. **(B)** Wild-type Caj1, along with the Patch 1, Patch 2 and Patch 1 and 2 Alanine mutants were incubated with liposomes containing PA to compare their relative lipid binding affinities. PCPEPA lipids (50:40:10 molar ratio) were reconstituted into liposomes and incubated with the indicated Caj1 mutants. Liposomes were isolated by ultracentrifugation (170,000 × *g*, 20 minutes) and washed with TSG. Co-sedimented proteins were analyzed by 12% SDS-PAGE and Coomassie Brilliant Blue staining (lanes 1-4). Aliquots of the mixtures prior to ultracentrifugation were loaded as controls (lanes 5-8).

We performed liposome sedimentation assays to assess the ability of the Caj1 mutants to bind PA (Figure 5B). Using PCPEPA liposomes reconstituted at a 50:40:10 molar ratio, wild-type Caj1 interacts with PA and sediments with the liposomes as expected (Figure 5B, lane 1). The “Patch 1” and “Patch 2” alanine mutants also display some affinity for PA (Figure 5B, lanes 2 and 3). Strikingly, however, PA binding is almost completely abolished with the “Patch 1+2” double mutant (Figure 5B, lane 4). This later result show that these two positively charged regions play a critical role in the interaction between Caj1 and PA.

## DISCUSSION

In our previous work, we combined the nanodisc with SILAC-based proteomics to identify the protein interactome of several *E. coli* integral membrane proteins ^14^. In the current study, we apply a similar quantitative proteomics strategy to identify novel phospholipid-interacting proteins, using the yeast *S. cerevisiae* as a model. The nanodisc is indeed an effective platform for the *in vitro* characterization of peripherally bound membrane proteins ^11, 13, 14, 30^, as it provides a lipid bilayer surface that is fully soluble in aqueous solution, and whose lipid composition can be precisely controlled ^11, 13, 14^.

First, we tested nanodiscs containing either yeast total lipid extracts, or nanodiscs having pure PE. In both cases we identified several Rab GTPases with elevated SILAC ratios. The Rab GTPase proteins play critical roles in regulating intracellular membrane trafficking in eukaryotes ^24, 31^, and they associate with lipid membranes in a GTP-dependent manner via a C-terminal polyisoprene lipid modification ^23, 24^. It is thought that the Rab GTPases preferentially bind to membranes containing lipid packing defects, likely due to the greater insertion of the prenyl motif into the lipid bilayer ^31^. Due to the conical shape of PE lipids, these packing defects may be especially present in our PE-rich nanodiscs, explaining why these Rab GTPases were enriched in our pulldown experiments.

Following these encouraging results, we next focused on identifying potential interactors of phosphatidic acid (PA). This low-abundance anionic phospholipid helps regulate essential cellular processes, such as vesicular trafficking, cell proliferation and response to external stimuli ^5, 7^. Analysis of previously identified PA-interacting proteins reveals that the only shared feature between PA-binding proteins correspond to clusters of positively charged amino acids such as lysine and arginine, which may be necessary for forming electrostatic interactions with membranes ^3, 5–7^. There is no other apparent sequence conservation for these clusters, rendering bioinformatic identification of novel PA-binding proteins difficult ^3, 4, 8^. Using nanodiscs enriched with phosphatidic acid, we here identify the J-domain containing protein Caj1. We subsequently validate the Caj1-PA interaction *in vitro* using nanodiscs and liposomes. Using liposome sedimentation assays, we show that the negative charge on the PA headgroup is necessary for mediating this interaction. The importance of the charge on the PA headgroup is underscored by our observation that Caj1-PA binding is influenced by environmental pH as well as by the lipid phosphatidylethanolamine (PE) ^5, 6, 27^. We also identify two clusters of positively charged residues (termed “Patch 1” and “Patch 2”) within the Caj1 primary sequence which are important for mediating interaction with PA.

What is the function of Caj1 *in vivo*? The biological role of Caj1 in yeast remains unclear, although recent work has shown that it may be involved in membrane protein quality control ^16,17^. Importantly, microscopy studies on *S. cerevisiae* expressing endogenous levels of GFP-tagged Caj1 also showed that Caj1 localizes to the plasma membrane ^17^. Our current study adds a novel dimension to this previous finding, as we show *in vitro* that the recruitment of Caj1 to membranes requires phosphatidic acid. An important avenue for future research will be to explore possible changes in membrane protein biogenesis in the presence of the “Patch 1+2” Caj1 mutant identified here, which does not bind phosphatidic acid. Such studies should be ideally carried out *in vivo* to reveal precisely how Caj1 is involved in membrane protein quality control.

Examining our proteomic dataset more broadly, we find that several phosphatidylinositol related proteins (Pdr16, Pik1, Inp53, and Pdr17) are also identified as binding partners of PA. We note that PA is an important synthetic precursor for different classes of phospholipids, including PI lipids ^3, 5^. Thus, it is not entirely surprising that PI kinases/phosphatases may be interactors of PA lipids. Indeed, a previous study has shown that PA is a specific activator of phosphatidylinositol 4-phosphate kinase ^8^. We also note that well-characterized phosphatidic-interacting proteins, such as Opi1 and Pah1, are not detected in our study. Opi1, a transcriptional repressor, binds phosphatidic acid in the endoplasmic reticulum (ER) membrane ^27, 32^. However, Opi1 recruitment to the ER membrane requires the tail-anchored protein Scs2, which was not present in our nanodisc preparations ^27, 32^. Thus, the absence of Opi1 from our dataset is not surprising. Similarly, binding phosphatidic acid phosphatase Pah1 to PA in the ER membrane requires the Nem1/Spo7 complex ^33^. Since these additional protein factors were not included in our nanodisc preparations, it is understandable that we did detect Pah1 in our experiments.

Altogether, our results highlight the applicability of our nanodisc/proteomic workflow for discovering and validating new and unexpected protein-phospholipid interactions, such as Caj1, a soluble Hsp40 that was initially predicted to be nuclear localized (https://www.uniprot.org/uniprot/F2QZU1). The role of Caj1 in membrane protein quality control remains ill-defined, but our current findings open up new avenues of research to further characterize this protein in the context of lipid specificity. More generally, the proteomic workflow presented in this study can be applied to the systematic identification of the peripheral membrane interactome as a function of the membrane lipid composition. Given the importance of the peripheral membrane proteins in mediating many essential cellular processes, we expect this to be an exciting avenue for future research.

## Acknowledgments

This work was supported by the Natural Sciences and Engineering Research Council of Canada (NSERC) to F.D. The mass spectrometry infrastructure was supported by the Canada Foundation for Innovation, and methods by the BC Knowledge Development Fund and the Genome Canada/Genome BC through the Pan-Canadian Proteomics Centre (214PRO) to L.J.F.

## Notes

### Competing Interest Statement

The authors have declared no competing interest.

